# Goat PRP14 (gPRP14) has xeno-antigenic properties and works as a vaccine in preclinical models of cancer

**DOI:** 10.1101/2022.07.04.498594

**Authors:** Saverio Minucci, Benedetta Bussolati, Paul E. Massa, Alessia Brossa, Roberto Ravasio, Mona Saadeldin, Genny Degani, Elli Papadimitriou, Amal Saadeldin, Antonio Salvaggio, Cristina Visintin, Giulia Rizzi, Stefano Ricagno, Laura Popolo, Maria Antonietta Vanoni, Pier Giuseppe Pelicci

## Abstract

We studied the activity of recombinant goat PRP14 (gPRP14), a member of the RID protein family, as a xeno-antigen in preclinical models of cancer. Antisera from rabbits and mice immunized with gPRP14 showed strong reactivity against several tumor cell types, which was absent towards normal cells: the tumor selectivity was related to surface and intra-cellular expression in tumor cells, and to an exclusively intra-cellular localization in normal cells. *In vitro*, binding to tumor cells was followed by cytotoxicity which could be rescued by the addition of excess soluble antigen. *In vivo*, an anti-tumor activity of immunization with gPRP14 was observed in murine syngeneic models of breast cancer and melanoma: the anti-tumor response was present when gPRP14 was administered in a preventive setting, and persisted upon repeated challenges with tumor cells in long-term survivor mice. Finally, we showed that both the humoral and T-cell mediated responses are needed for the optimal anti-tumor effect in the murine melanoma model. Thus, we have performed an initial characterization of gPRP14 as a cancer vaccine, which -given the potential wide range of tumor cells positive for the antigen-appears as a promising, novel immunotherapy.

## Introduction

The UK114 family of proteins (also referred as YjgF or YERO57c7) is highly conserved in evolution. Though different putative functions have been described for single members, a common enzymatic metabolic activity (reactive intermediate/imino deiminase) has been recently identified as a key biochemical feature of all family members (1).

UK114 was originally identified in the perchloric acid extract of mammalian liver and shown to possess anti-cancer activity (2). In particular, antisera obtained from rabbits or mice immunized with the goat extractive UK114 were shown to target selectively human and rodent cancer cells (2-3).

The native UK114 protein is cytoplasmic in normal cells and both cytoplasmic and membrane-associated in transformed cells (2-3). Indeed, monoclonal antibodies raised against extractive UK114 bind to and mediate cytotoxicity against several cancer cell lines (of gastric, breast, colonic and neural origin), but not the corresponding normal cells, lymphocytes or monocytes (3-4). Extractive UK114 has been evaluated in preliminary clinical studies, which demonstrated low toxicity and some efficacy in patients with different tumor types (5).

In order to eliminate potential biases due to the extractive origin of the protein, we produced recombinant UK114. Crystallographic studies revealed a 3D oligomeric conformation (6) which occurs mainly as a trimer in solution (7). We produced the His-tagged recombinant protein from goat (hence, gPRP14), which can be purified as a highly stable homotrimer (8). The availability of the recombinant protein allows deeper investigations of its immunogenic properties. In the present study, we show that mouse and rabbit sera raised against gPRP14 match the properties previously obtained with sera against the extractive protein. In addition, experimental evidence of the effect of vaccination with PRP14 in mouse tumour models allows to trace prospects for exploitation of this tumour-associated xeno-antigen.

## Materials and Methods

### Recombinant goat PRP14 (gPRP14) and cells

gPRP14 was produced and purified as described (8). TUBO cells (mouse mammary carcinoma cells) were kindly provided by Federica Cavallo (University of Turin). HT-29 (human colon carcinomas) and HK-2 (human renal epithelial cells) cell lines were obtained from ATCC, and cultured in DMEM high glucose, supplemented with 10% FCS and antibiotics. B16.F10 mouse melanoma cells and primary tumor cells (MLL-AF9 leukemic cells) were cultured in RPMI supplemented with 20% FCS, antibiotics, sodium pyruvate [1mM] and non-essential amino acids [0.1mM]. K562 human chronic myelogenous leukemia and CT-26 murine colorectal carcinoma cells were cultured in RPMI supplemented with 10% FCS and antibiotics. Wild type and MMTV-primary mammospheres were derived and cultured as previously described from primary breast tissue obtained from Balb/c female mice with palpable MMTV-tumor masses, or from tumor-free littermates. (9).

### ELISA assays

The presence of antibodies against gPRP14 in sera derived from immunized animals was evaluated by ELISA. Briefly, purified gPRP14 was left to bind overnight in 96-well plates (2μg /well) in 100μl of Sodium Bicarbonate buffer followed by blocking with PBS-5% BSA (Sigma Aldrich) for 1 hour. 100 μl of serum from immunized animals (rabbits and mice), or pre-immunization serum as control, diluted in PBS-5% BSA (dilution range: 0.01% - 1%), was added to the wells and incubated for one hour at room temperature. Four washes with 400 μl of PBS-0.5% Tween-20 followed. The appropriate secondary antibodies (anti-Rabbit or anti-Mouse IgG, HRP-conjugated from Dako) were added at a dilution 1:10.000 in PBS-5%BSA and incubated at room temperature for one hour, followed by four washes with PBS-1% BSA. Finally, 100 μl/well of TMB solution (Sigma Aldrich) was added according to manufacturer’s instructions. The reaction was stopped, and the signal was evaluated by absorbance using a microplate reader (Bio-Rad). Data are expressed as mean ± SD of the media absorbance of different experiments performed in triplicates.

### Western blot

Purified proteins were separated by SDS-PAGE (12 % acrylamide) gel and transfered onto a nitrocellulose membrane. Rabbit antiserum anti gPRP14 was used at diluition of 1:10,000 and incubated 1h at room temperature. Anti-rabbit conjugated with Alexa 488 (Thermofisher) was used as secondary antibody and the signal was detected in a ChemiDoc MP (BioRad).

### Cytotoxicity assays

For cytotoxity experiments, we used different cell lines. Briefly, HT-29 cells were plated in 24-well plates (10.000 cells/well) in DMEM high glucose-10% FCS. As soon as they attached, new medium (DMEM low glucose from Lonza, supplemented with 1% rabbit complement sera from Sigma-Aldrich) containing 10% sera derived from immunized or not with gPRP14 rabbits or mice was added. Forthy-eight hours later, Annexin V apoptosis assay was performed using the Muse™ Annexin V & Dead Cell Kit (Millipore), according to the manufacturer’s recommendations. Alternatively, TUBO cells were plated in 96-well plates (3.000 cells/well) in DMEM high glucose-20% FCS. After 24 hours, the medium was substituted with new DMEM low glucose, containing sera from immunized rabbits (range from 0,5 to 10%) and a remaining amount of FCS to reach 20%. As control, sera from rabbits before immunization was also used. After 72 hours, the vabilitiy of TUBO cells was evaluated by MTT assay, according to the manufacturer’s protocol.

### Flow Cytometry on cells and mammospheres

For cell lines/primary cells, 500,000 live cells were harvested and co-incubated for 30 minutes at 4 degrees in the presence of 2 microliters of either normal rabbit pre-immune plasma, or post-immune anti gPRP14 plasma (or in the absence of plasma as a further control). Cells were pelleted and washed with PBS twice and then co-incubated in detection antibody (donkey anti rabbit IgG-AF488: Abcam[Ab150093])) diluted 1:5000 in PBS and incubated for 15 minutes at 4 degrees. Cells were washed and immediately acquired on a Becton-Dickinson Celesta flow cytometer.

### In vivo studies

All animal studies were conducted in accordance with the national guidelines and regulations and were approved by the Italian Ministry of Health (Protocol Number: 959/2018-PR; 613/2020-PR; 338/2022-PR). For rabbit immunization, New Zealand white rabbits (12 weeks old) were injected subcutaneously with gPRP14 (6 mg/rabbit) at days 1, 8, 22 and 37. gPRP14 was administered in 50% Freund’s Adjuvant. At the end of the immunization period, a final heart bleeding was performed.

For experiments in the murine breast cancer syngeneic model (murine breast cancer TUBO cells), 6-8 weeks old male Balb-C mice were used. In the therapeutic protocol, mice were first subcutaneously injected with 100×10^3^ TUBO cells, and gPRP14 protein subcutaneous injection (10 µg/mouse) started one week after tumor implantation. gPRP14 was diluted in 100 μl physiologic solution, mixed with 100 μl of Freund’s Adjuvant (Sigma-Aldrich), using a 1-ml syringe with 26-gauge needle. In vaccination experiments, mice were injected once/week for four (or eight) consecutive weeks with gPRP14 (10 µg/mouse). Control groups received physiologic solution mixed with Freund’s Adjuvant only. After the immunization period, TUBO cells were injected as described above. 2 plugs/mouse were performed. In both settings, three weeks after tumor cell implantation mice were sacrificed and sera were harvested for testing the reactivity against gPRP14 by ELISA (see legends to the figures). In parallel, tumor volume was measured. Mice were sacrificed when tumor larger diameter was more than 1 cm. Survival rate was therefore calculated. For all the *in vivo* experiments, tumor volume was measured with a caliper and calculated using the formula: v= (L × l^2^)/2, where “L” indicates the larger diameter and “l” indicates the smaller one.

In the murine melanoma syngeneic model, C57BL/6J 6-8 week old female mice were bought from Charles River. Purified proteins (ovalbumin as control, gPRP14) were resuspended in PBS at the concentration of 1 or 10 µg/50 µl, and added to 50 µl of complete Freund’s adjuvant, they were resuspended in emulsion and injected intradermically into the rear flanks of animals (100 µl per animal per dose). Doses were administered as indicated in the legends to individual figures.

Vaccinated or control animals (n=5-6 animals per group) were implanted subcutaneously in the contralateral (non-immunized) flanks with B16F10 cells (either 10,000 or 250,000 cells) in 100 µl of PBS. Tumor growth was monitored daily and dimensions were measured by calipers. When tumor area exceeded 150mm^2^, animals were sacrificed humanely. Kaplan-Meier survival curves were generated using Prism statistical software to show time-to-sacrifice and significance was determined by log-rank and Mantel-Cox tests.

### T cell adoptive transfer

Mice were inoculated three times (every 10 days) with 10µg of either ovalbumin or gPRP14. 2 weeks after the last vaccination, animals were sacrificed and spleens were harvested. CD3-positive T cells were purified using the Pan T Cell Isolation kit (Miltenyi). 50 million CD3 cells from control animals were stimulated with Dynabead CD3/CD28 activator beads in an antigen-independent manner, while CD3+ cells from vaccinated animals were put in co-culture with primary bone marrow-derived macrophages (1:1 ratio) that had been previously loaded with the protein used to immunize mice. T cells were grown in medium supplemented with murine IL-2 (20ng/ml, Peprotech) for 4 days. On day 1 of the T cell culture, control 10 week-old recipient C57BL/6J female mice were non-lethally irradiated to diminish circulating T cells, while on day 2 animals were transplanted with 10,000 B16F10 cells. 2 days later, stimulated T cells were injected into transplanted animals (1.5×10^6^ CD3+ cells per animal) and tumor growth was monitored as described above (n=12-14 mice per group).

### Post-immune serum treatment

Mice were inoculated three times (every 10 days) with 10µg of either ovalbumin or gPRP14. 2 weeks after the last vaccination, animals were sacrificed and heart blood was collected, pooled, and allowed to clot for 45 minutes on ice. Post-clotting sera were collected and aliquoted for subsequent use. Untreated 10 week-old C57BL/6J female animals were transplanted with 10,000 B16F10 cells and beginning from day 2 post-transplant animals were treated with post-immune sera twice per week for up to 4 weeks. Animals were treated either intraperitoneally with 300 microliters of serum, or 20 microliters intradermically by Hamilton syringe adjacent to the site of transplant. Tumor growth was monitored and animals sacrificed as described above (n=4 mice per group).

## Results

### gPRP14 triggers an immune response against tumor cells in vitro

The biochemical properties of the purified gPRP14 have been described previously (8). To investigate its antigenicity, the purified protein was injected into rabbits and the corresponding antisera analysed by ELISA against immobilized gPRP14 (supplementary figure 1A) and western blotting against the SDS-denatured protein (supplementary figure 1B). As expected, the antisera were highly reactive against gPRP14 in both assays.

FACS-analyses of reactivity of the anti-gPRP14 rabbit antisera plasma against several cancer cell lines showed consistent positive-staining of both human (K562 erythroleukemia; CT26 colorectal) and murine (B16.F10 melanoma; MML-AF9 AML) cell lines (fig. 1A). To investigate specificity of binding to cancer cells, we then tested FACS-reactivity against normal versus transformed primary epithelial mammary cells, obtained from wild-type or MMTV-ErbB2 transgenic mice, respectively, and grown in vitro as mammospheres (see methods). Upon permeabilization of cell membranes, both normal and transformed cells showed strong and comparable reactivity against the anti-gPRP14 antisera. Strikingly, however, only tumor cells tested positive when the antisera was incubated on intact cells, which expose only membrane antigens (fig 1B). These results confirm previous studies with the extractive UK114 showing cytoplasmic localization of gPRP14 in normal cells, cytoplasmic and membrane-associated in cancer cells (3).

**Figure 1.**
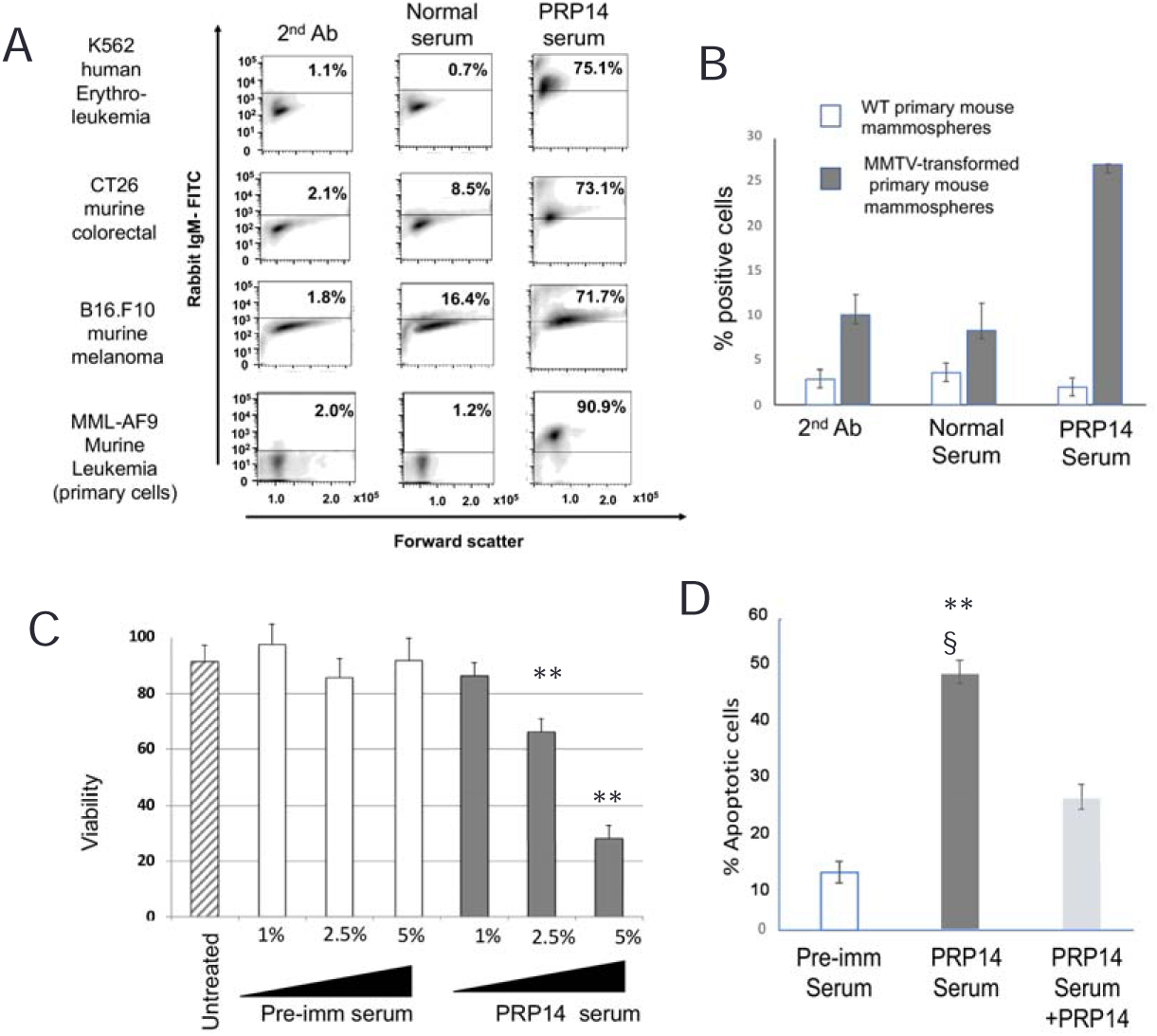
gPRP14 antisera react with transformed cells and mediates tumor cell cytotoxicity. A) Flow cytometric analysis of intact, live cells incubated with normal pre-immune and gPRP14 post-immune rabbit plasma; B) Flow cytometric positivity of wild-type or MMTV-transgenic mice mammary tissue derived mammospheres incubated with normal pre-immune and GPRP14 post-immune rabbit serum. C) Dose dependent cytotoxic activity of rabbit PRP serum but not pre-immune rabbit serum (pre-imm), on TUBO murine breast cancer cells as assessed by MTT assay. Data are mean + SD of five different experiments. ANOVA with Bonferroni’s post hoc test: **=p<0.001 vs untreated. D) Percentage of apoptotic HT29 colon cancer cells treated for 48h with preimmune serum, rabbit PRP serum or rabbit PRP serum pre-incubated with recombinant gPRP14 (PRP serum+PRP antigen). Data are expressed as mean ± SD of the percentage of Annexin V^+^ positive cells (n=3). ANOVA with Bonferroni’s post hoc test : **=p<0.001 vs pre-imm; §=p<0.05 vs PRP serum+PRP antigen.

We then evaluated the in vitro cytotoxic activity of the anti-gPRP14 antisera. Tumor or normal cells were incubated with the anti-gPRP14 antisera or control serum and viability analysed using the MTT assay. A dose-dependent effect on cell viability was observed against murine breast (TUBO cells, expressing a constitutively active rat NeuT - Her-2 homologue) and melanoma (B16.F10) tumor cells (fig. 1 C, supplementary fig 2A), and against the human HT29 colon cancer cells (supplementary fig. 2B), which was reverted by absorption of the antisera with the recombinant antigen (fig. 1D and supplementary fig. 2C). The anti-gPRP14 antisera had no effect on HK-2 normal renal epithelial cells, which do not express the gPRP14 antigen (supplementary fig. 2B).

### gPRP-14 vaccination shows an anti-tumor activity in syngeneic murine models of melanoma and breast cancer

We then tested the effects of recombinant gPRP-14 protein vaccination *in vivo*, using syngeneic models of murine melanoma (B16-F10 cells) and breast cancer (TUBO cells). B16-F10 and TUBO tumors were modelled in Balb/c and C57BL/6J mice, respectively, thus allowing analyses of strain-dependency.

In initial experiments, we measured the effect of gPRP-14 vaccination in a “therapeutic” setting, e.g. administering the antigen to mice already harboring growing tumors. We did not detect, however, a strong anti-tumor effect of gPRP-14 injection in either tumor models, although for breast cancer we observed a trend towards an increased survival, with one long-term survivor in the vaccinated mice cohort (Supplementary figure 3).

We then tested the effect of gPRP-14 in a “preventive” setting, administering gPRP-14 before inoculation of the mice with tumor cells, to allow generation of immune reactivity prior to tumor cell injection. Mice were subcutaneously injected once/week for four consecutive weeks with 25µg/mouse of recombinant gPRP-14 antigen (or control), and then injected with TUBO tumor cells. Tumor growth was monitored for the subsequent three weeks (figure 2A). Strikingly, gPRP-14 administration almost completely prevented tumor expansion, leading to a marked reduction of tumor size at the time of mouse sacrifice (figure 2C-D). Analyses of anti-gPRP14 antibodies by ELISA showed increased reactivity at the time of injection of tumor cells, which increased further at sacrifice (figure 2B).

**Figure 2.**
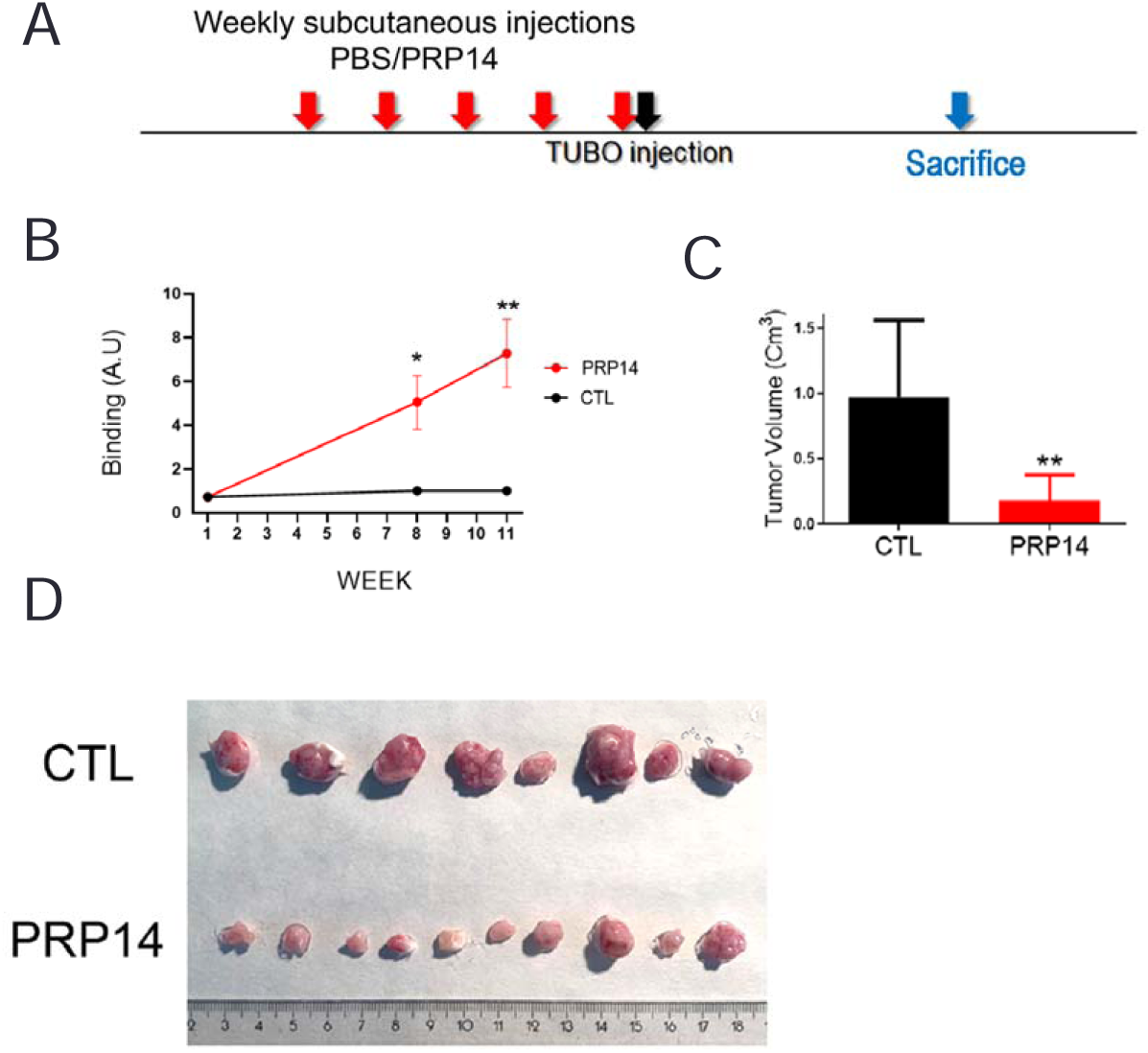
gPRP14 vaccination delays murine breast cancer growth. A) Scheme of the protocol. Balb-c mice were subcutaneously injected once/week for four consecutive weeks with 1mg/kg gPRP14 (25 µg/mouse) in Freund’s Adjuvant. Control group (CTL) received PBS in Freund’s Adjuvant only. 100×10^3^ TUBO cells were then injected subcutaneously, and animals sacrified after 3 weeks. B) ELISA test showing the binding of murine sera from CTL or gPRP14-immunized mice at different times upon injection of TUBO cells. C) Assessment of tumor volume in CTL and gPRP14-immunized animals at sacrifice. D) Images of tumor plugs recoved after sacrifice. N=8 animals/group. **p<0.01 vs CTL.

Similar results were obtained with the melanoma B16F10 model. Injection of gPRP14 at 1µg/mouse delayed tumor growth and slightly increased survival of tumor-bearing mice (supplementary figure 4). Injection of gPRP14 at 10 µg/mouse (same schedule as in figure 3A) induced a markedly stronger effect on tumor growth (figure 3B) and survival (figure 3C), with 50% mice surviving for more than 2 months from the inoculation of tumor cells. Injection of same amounts of ovalbumin (as control) showed no effect on tumor growth or mouse survival (figure 3B-C).

**Figure 3.**
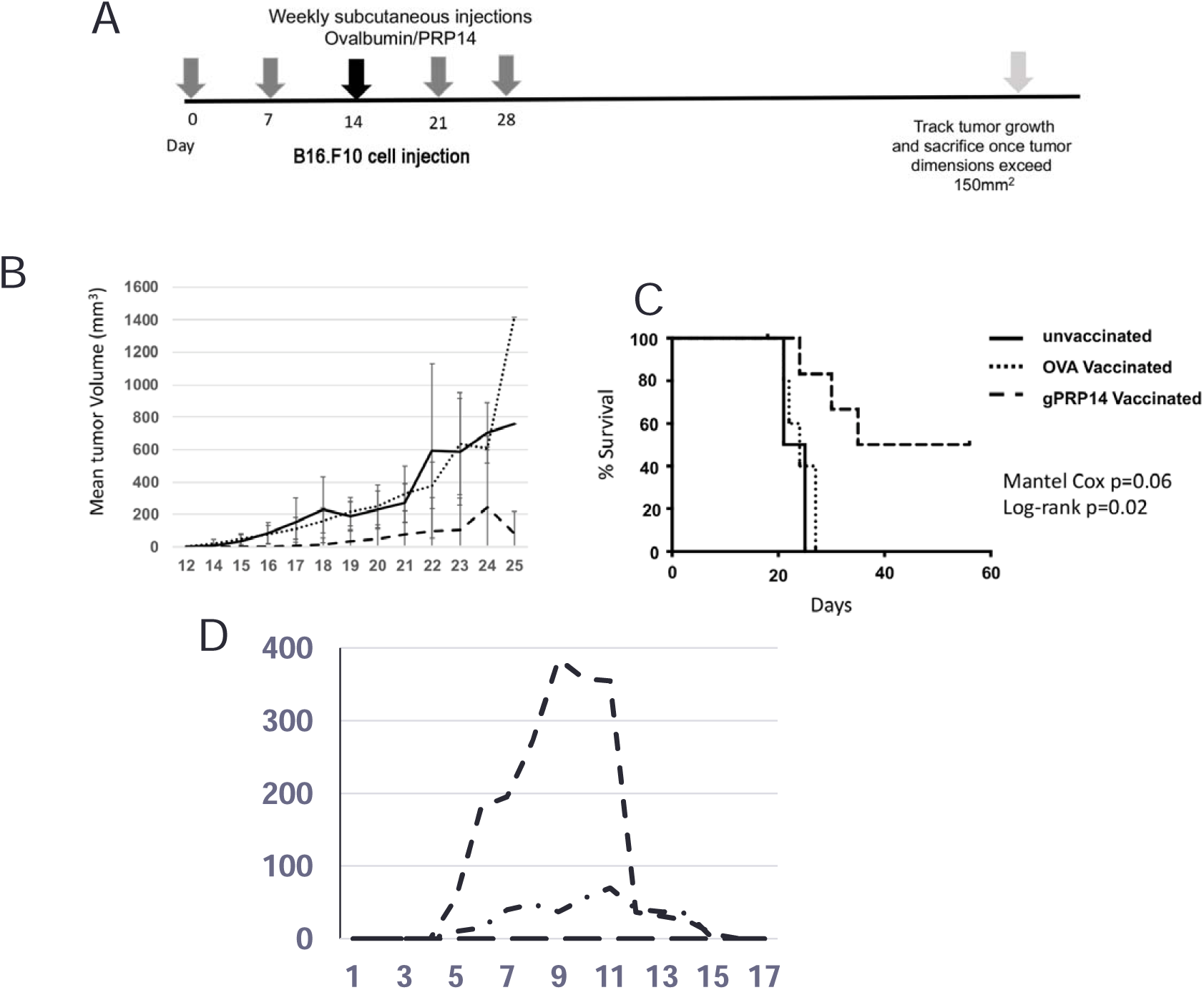
Immunization with gPRP14 protects mice against B16F10 mouse melanoma engraftment and induces long-lasting anti-tumor activity. A) Schedule of immunization/treatment. B) Mean tumor volume measured post-engraftment of B16F10 in gPRP14-immunized (dashed line), Ovalbumin-immunized (dotted line) or non-immunized (solid line) mice. C) Kaplan-Meier survival analysis of mice engrafted with B16F10 melanoma cells, and treated as indicated. D) Tumor volume growth curves of individual, long-term gPRP14-immunized survivor mice post-rechallenge with B16F10 tumor cells.

To evaluate the establishment of protective, long-term immunity in the vaccinated mice, we re-challenged 3 survivor mice of the gPRP14 vaccinated group with B16.F10 melanoma cells, without further antigen administration. 1/3 mice did not show any signs of tumor growth, while the remaining 2/3 mice developed measurable tumors, which however then regressed (Figure 3D). ELISA testing for reactivity against gPRP14 in the mouse plasma showed comparable reactivity in mice immediately after immunization and at the time of the re-challenge with tumor cells, supporting the view that the protective anti-tumor effect is due to establishment of long-term immunological protection (supplementary figure 5).

### Both humoral and T-cell mediated immunity are involved in the immune response triggered by the gPRP14 antigen

As shown above, immunization with gPRP14 triggers the production of antibodies with in vitro anti-tumor cytotoxic activity. We therefore investigated in the murine melanoma model whether sera from immunized mice can transfer anti-tumor activity. Notably, intra-peritoneal injection of serum from gPRP14-injected mice delayed tumor growth and prolonged survival of melanoma-bearing mice, as compared to the control serum of mice injected with ovalbumin (figure 4A and 4C). We also tested whether route of administration (systemic versus intradermic/intratumoral) influence the effect of the anti-gPRP14 antisera, as shown for other cancer immunotherapies (10). Results showed that injection of the anti-gPRP14 antiserum adjacent to the tumor had better therapeutic efficacy as compared to the systemic administration (figure 4B and 4D). Thus, we confirmed a critical role for humoral immunity triggered by the xenogenic gPRP14 antigen in the anti-tumor activity.

**Figure 4.**
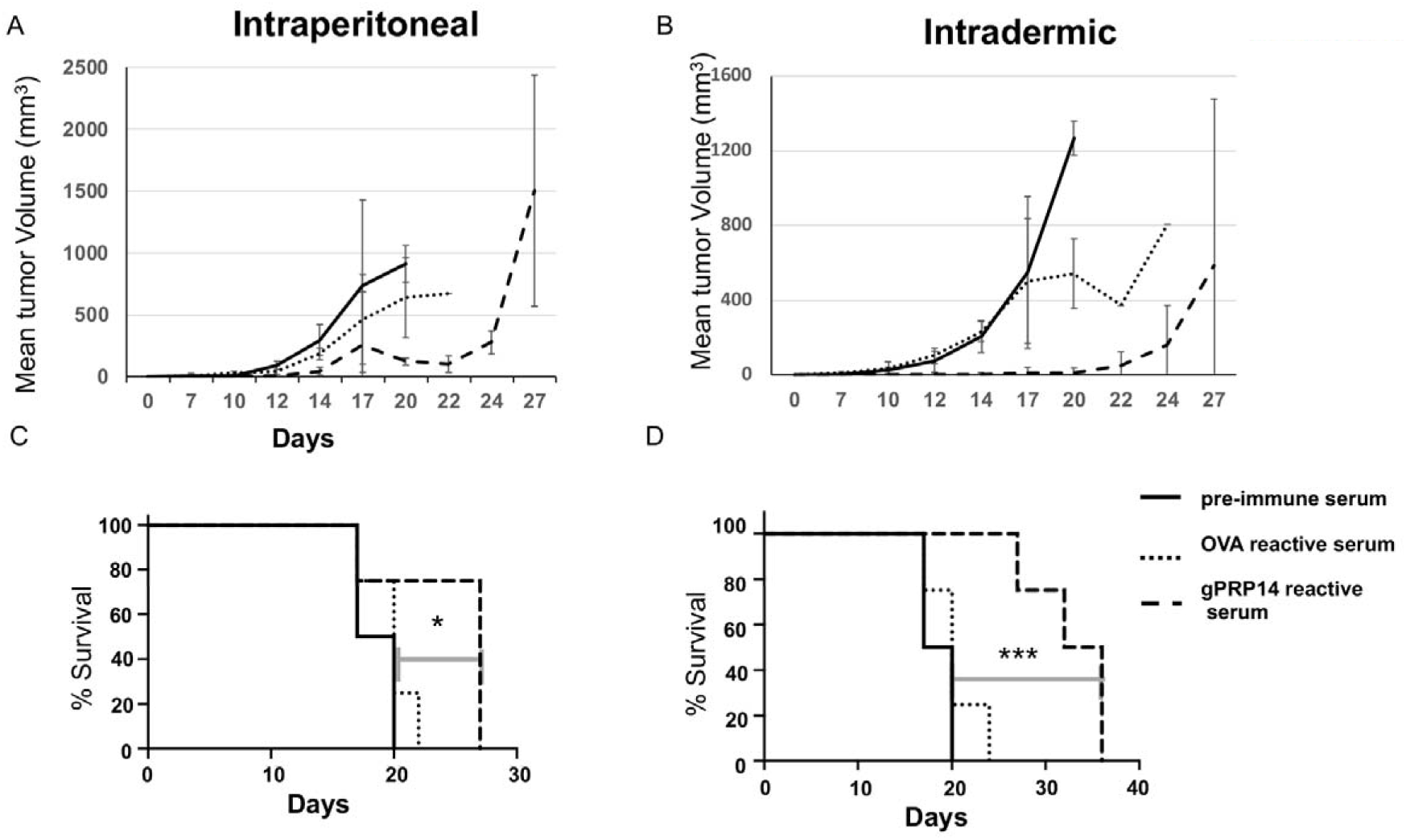
gPRP14 reactive sera protects against B16F10 tumor growth. Mean tumor volume growth curves of mice treated intraperitoneally (**A**) or intradermically (**B**) with non-reactive pre-immune sera (solid line), ovalbumin-reactive sera (dotted line) or gPRP14 reactive sera (dashed line); Kaplan-Meier survival curve analysis of intraperitoneal (**C**) or intradermic (**D**) treatment, as described above.

Antisera treatment, however, did not recapitulate entirely the effect observed by antigen administration (compare figures 4 and 3). Though the different experimental contexts may explain this difference, we investigated whether this might also be due to a contribution of T cells to the anti-tumor activity. We therefore administered CD3+ T cells derived from mice immunized with gPRP14 (or ovalbumin as control) to mice previously injected with B16F10 melanoma cells. Since purified T cells were further exposed to gPRP14 (or ovalbumin as control: see methods) *in vitro*, we also tested purified CD3+ T cells stimulated aspecifically *ex vivo* by CD23/CD28. Importantly, only T cells exposed to the gPRP14 antigen strongly delayed melanoma tumor growth, and prolonged survival (figure 5 A-B). Even in this case, however, the effect observed did not fully recapitulate the vaccination effect (compare figures 3 and 5), suggesting that both humoral and T-cell immunity contribute to the anti-tumor immune response triggered by the gPRP14 antigen administered in vaccination settings.

**Figure 5.**
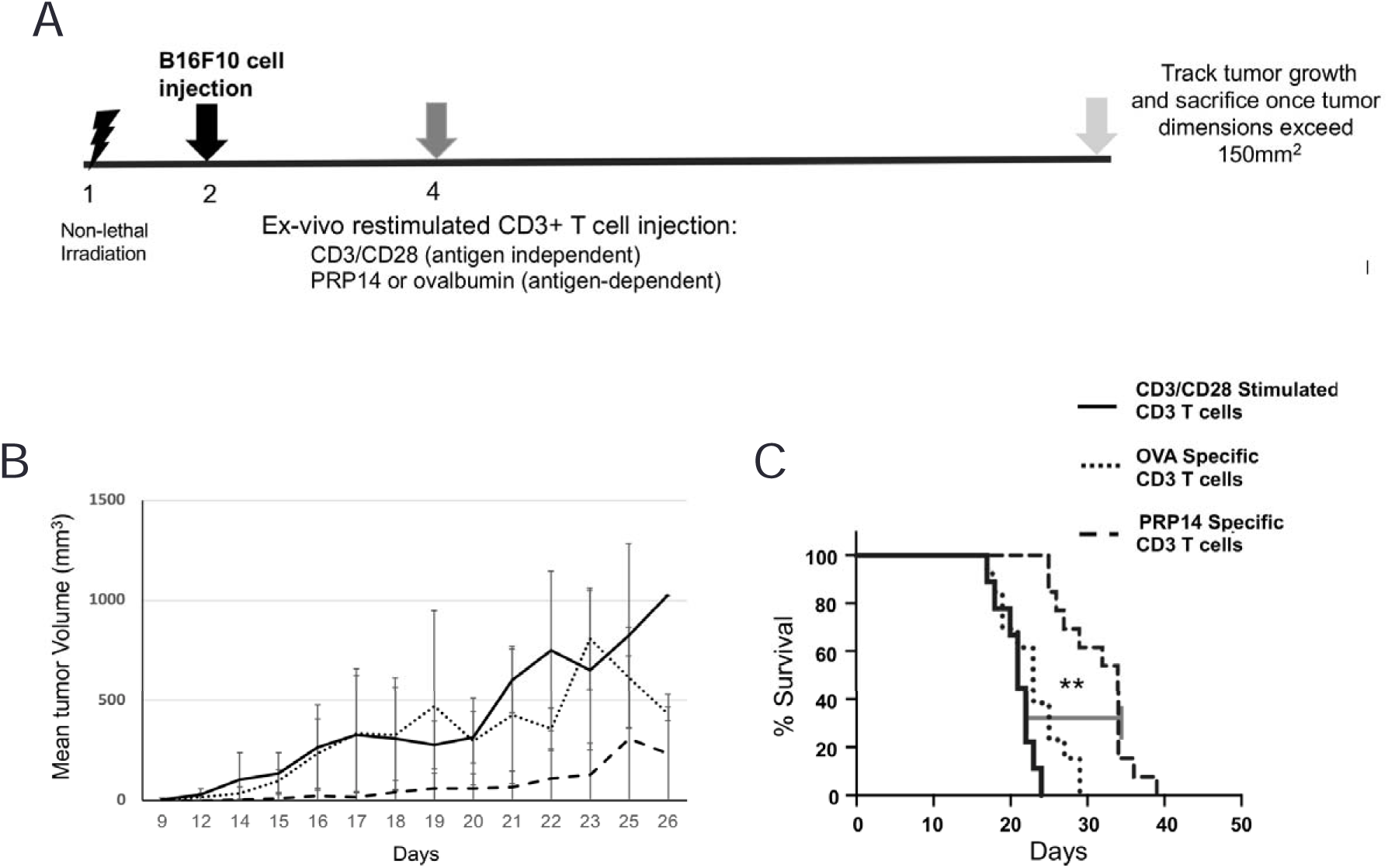
gPRP14-specific T lymphocytes transfer anti-tumor immunity into unvaccinated B16F10 bearing recipients. A) Schedule of T cell treatment and tumor implantation. B) Mean Tumor volume growth curves of mice treated with in vitro expanded antigen aspecific CD3/CD28 stimulated (solid line), Ovalbumin specific/stimulated (dotted line) or gPRP14 specific/stimulated (dashed line) CD3+ T lymphocytes. C) Kaplan-Meier survival curves of T-cell treated animals.

## Discussion

Recent advances in cancer immunotherapy have dramatically changed the natural history of many tumor types. The use of monoclonal antibodies against tumor targets, the introduction of cellular (CAR-T) therapies and of immune-checkpoint inhibitors has demonstrated that it is possible to re-establish through different approaches a potent immune response against tumor cells, which in some cases may lead to complete eradication of the disease. However, even in the face of these significant advances, a significant fraction of cancer patients is unresponsive to immunotherapies, or relapse after treatments (11).

Among the different cancer immunotherapy approaches, patient vaccination against tumor-associated antigens (either mutated or wild-type proteins expressed by tumor cells) is lagging behind, despite the strong rationale and encouraging clinical results, mainly due to concerns on its feasibility. A core hurdle is the observation that most mutated tumor-associated antigens are “private” and expressed in a patient-specific manner, thus imposing the production of many antigens for vaccination. An alternative approach is the usage of non-mutated antigens with altered expression patterns in tumor cells. In this case, however, antigenicity may be poor, due to biological restraints limiting the induction of auto-antibody formation, thus resulting in a weak immune response and negligible anti-tumor activity. To circumvent these limts, a xeno-antigenic approach has been proposed to overcome tolerance, providing antigens sufficiently distinct from those “known” to the host immune system (12). While this approach has been validated both at the preclinical and clinical level, it is currently not widely pursued, most likely due to the success of the other immunotherapic strategies, as summarized above (12).

Here, we report a proof-of-concept preclinical study of the anti-tumor activity of the gPRP14 xeno-antigen. The goat protein is highly similar to its rabbit and murine (and human) orthologues. We observed a strong anti-tumor activity of gPRP14 on both tumor-models investigated, the murine breast cancer TUBO cell line and the B16F10 murine melanoma cell line. Strikingly, the immune response showed high selectivity against tumor cells, which may be the only exposing the antigen, due to its membrane localization, as compared to normal cells.

Since PRP14 possess a detoxifying enzymatic activity, expression on the cell membrane of tumor cells might be related to a defence mechanism of the highly proliferating cancer cells to the toxic metabolic products accumulating in the microenvironment. Indeed, RidA proteins have been proposed to counteract the metabolic stress caused by the increased metabolism of serine, which is typically observed in highly proliferating cancer cells (13).

We showed that gPRP14 is clinically active in cancer models in either the Balb/c or C57BL/6 murine strains. Since these strains show extensive morpho-functional differences in their immune system, these results suggest that adaptive responses to the xeno-antigen are similarly effective in different contexts, and that this approach may be extended to several tumor types which show surface immune-reactivity, thus defining surface staining as a candidate biomarker for patient stratification.

The anti-tumor efficacy of gPRP14 was significantly stronger upon challenge in a “vaccination” setting, where gPRP14 is administered before transplantation of cancer cells, as compared to a “therapeutic” setting, where antigen administration followed grafting of tumor cells. We surmise that this difference is linked to the fact that experimental tumor grafts have a very rapid growth, thus preventing the establishment of an efficient immune response de novo. This may possibly not be a limitation in the human setting, since the rate of growth of spontaneous human cancers is considerably slower than in our experimental models. Regardless, our re-challenge studies also validate a “vaccination-like” approach, following tumor eradication (by surgery or other treatments), to prevent disease relapse. Further studies in animal models may lead to optimization of clinical strategies.

The observed anti-tumor effect was dependent on both host humoral and cell-mediated immune responses, as revealed by adaptive transfer experiments. In-depth analyses of immune responses following immunization is ongoing in our laboratories to address mechanistically the different immune components of the response observed. Our results also suggest that a recently described, direct cytotoxic effect of a mushroom member of the RidA family on cancer cell lines is not playing a key role in our setting (14).

Finally, it remains to be tested whether the observed immune response is influenced by active immune checkpoints, thus providing an approach to enhance efficacy by combining the xeno-antigen approach with checkpoint inhibitors (anti PD1/PD-L1, anti-CTLA4).

Our studies stem from pioneering work performed with an extractive form of the presently used gPRP14 (UK114). The extractive UK114 was also used in initial clinical studies, which showed lack of toxicity and encouraging signs of efficacy. Recombinant gPRP14 is easy to produce in large amounts, high purity and with qualitative standards which cannot be easily reached with the extractive form, thus providing a strong advantage for its clinical development.

## Acknowledgements

We thank the members of our groups for useful discussions. We thank: Professor Alberto Bartorelli -who discovered UK114-for scientific discussions and continuous feedback; Professor Giovanni Bussolati -who achieved key findings on UK114-, Professors Alberto Panerai and Paola Sacerdote, for scientific discussions and critical evaluation of the manuscript. We acknowledge the generous help of Fondazione Sisini. This work was partially supported by the Italian Ministry of Health with Ricerca Corrente and 5×1000 funds.

