## Supplemental figures for "Goat PRP14 (gPRP14) has xeno-antigenic properties and works as a vaccine in preclinical models of cancer"

### Supplementary figures

A

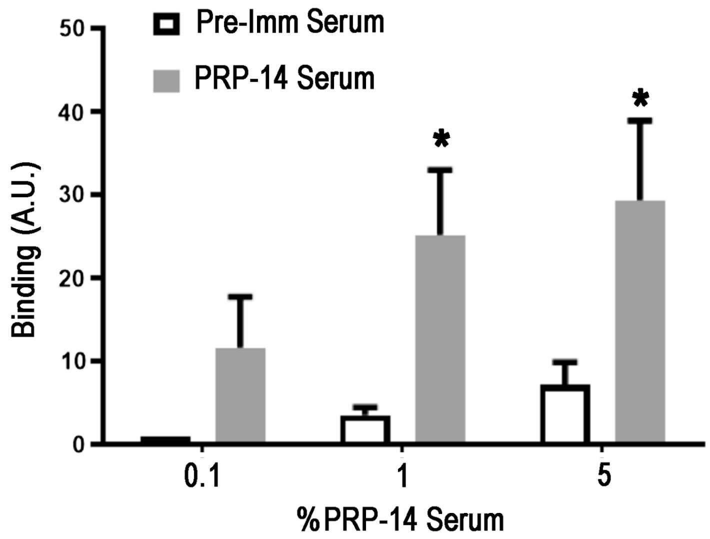


B

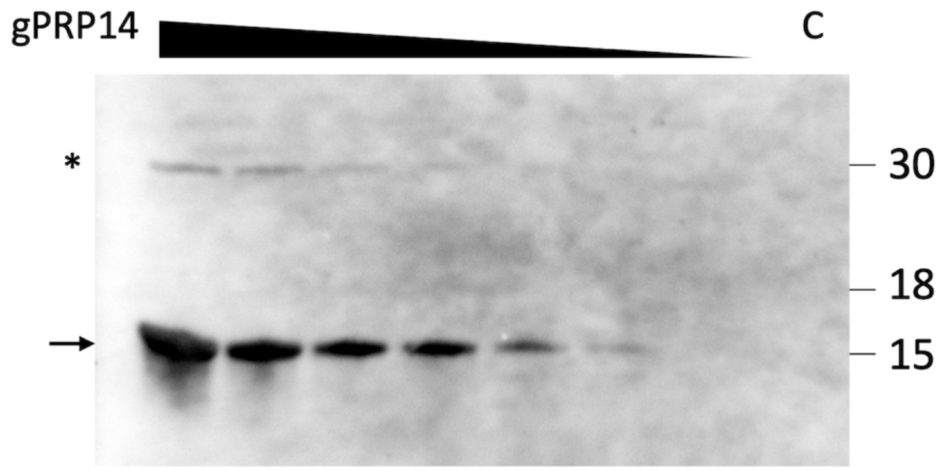


Supplementary Figure 1. Biochemical characterization of rabbit anti-gPRP14 sera. A) Dose-dependent anti-PRP14 reactivity was assessed as binding to the recombinant gPRP14 antigen by ELISA. Pre-immune serum (Pre-Imm) was used as control. Data are mean +SD of 5 different experiments. T-test: *p<0.05. B) Decreasing concentrations of purified gPRP14 (1, 0.50, 0.25, 0.10, 0.05, 0.02 and 0.001 μg) were analysed by Western Blot; 4μg of beta-2-microglobulin were used as control (lane C). Monomeric gPRP14 is indicated by the arrow and the dimeric gPRP14 by the asterisk.

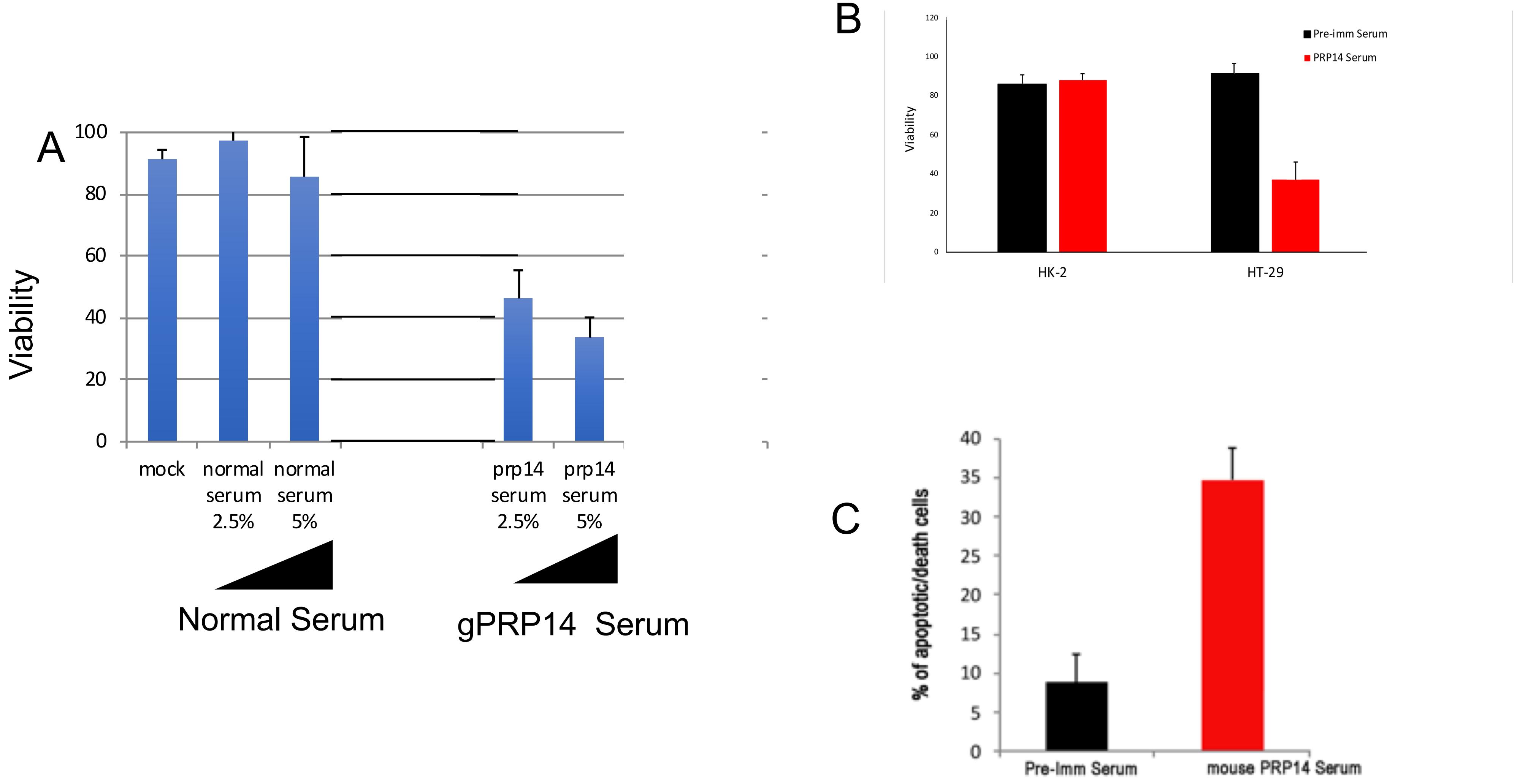


Supplementary Figure 2. Cytotoxic activity of gPRP14 antiserum on human and murine tumor cells. A) Dose dependent cytotoxic activity of mouse PRP serum but not of normal murine serum, on B16F10 murine melanoma cells, as assessed by MTT assay. Data are mean + SD of five different experiments. B) Selective effect of gPRP-14 antiserum on HT29 colon cancer cells but not on HK-2 normal renal epithelial cells, as assessed by MTT assay. Data are mean $\pm$SD of 3 different experiments. **C**. Percentage of apoptotic TUBO murine breast cancer cells after treatment with mouse gPRP14 antiserum or mouse pre-immune (Pre-Imm) serum. Data are expressed as mean ± SD of the percentage of Annexin $V^+$ positive cells (n=3). T-test: *p<0.05 vs Pre-Imm.

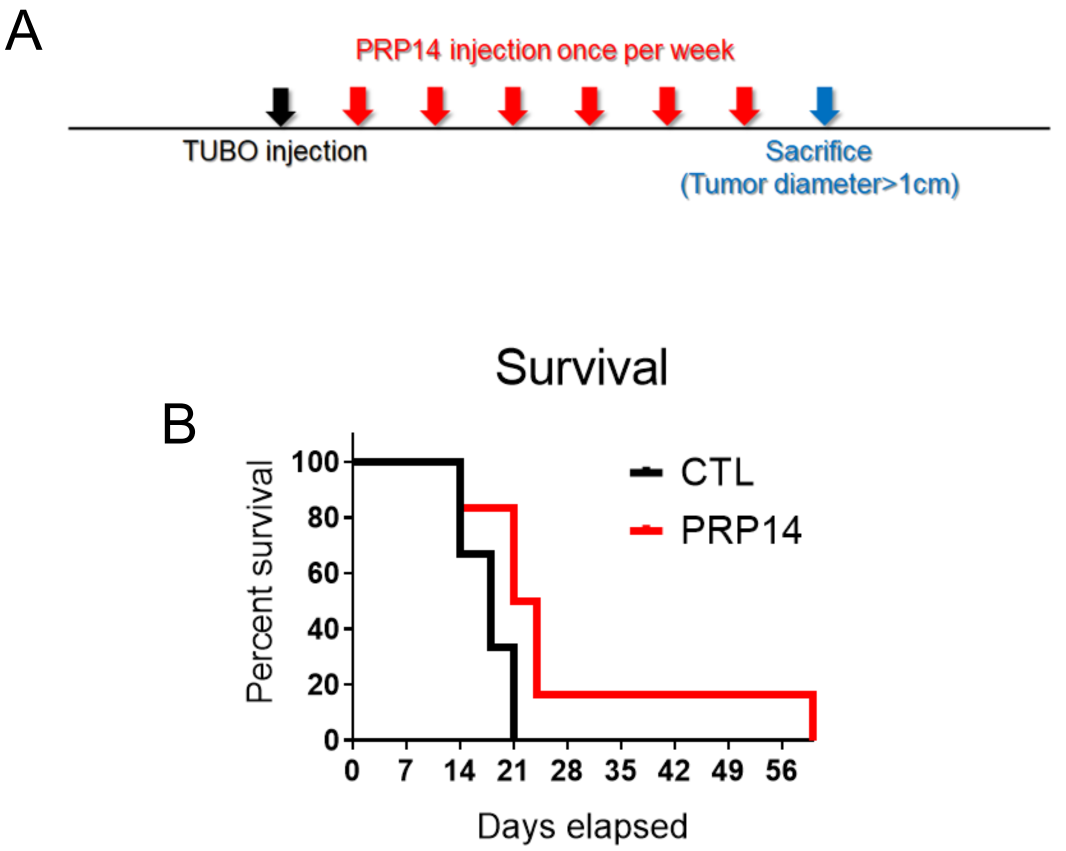


Supplementary Figure 3. In vivo effect of PRP14 vaccination on murine breast tumor growth in a therapeutic setting. A) Scheme of the protocol. Balb-c mice were subcutaneously injected with $100x10^3$ TUBO cells and subsequently treated once/week with 1mg/kg (25 µg/mouse) gPRP14 in Freund's Adjuvant. Control group (CTL) received PBS in Freund's Adjuvant only. Mice were sacrificed when tumor diameter was more than 1 cm in any of the diameters. B) Survival rate showing mice survival after treatment with gPRP14 or PBS (CTL). N=6 animals/group.

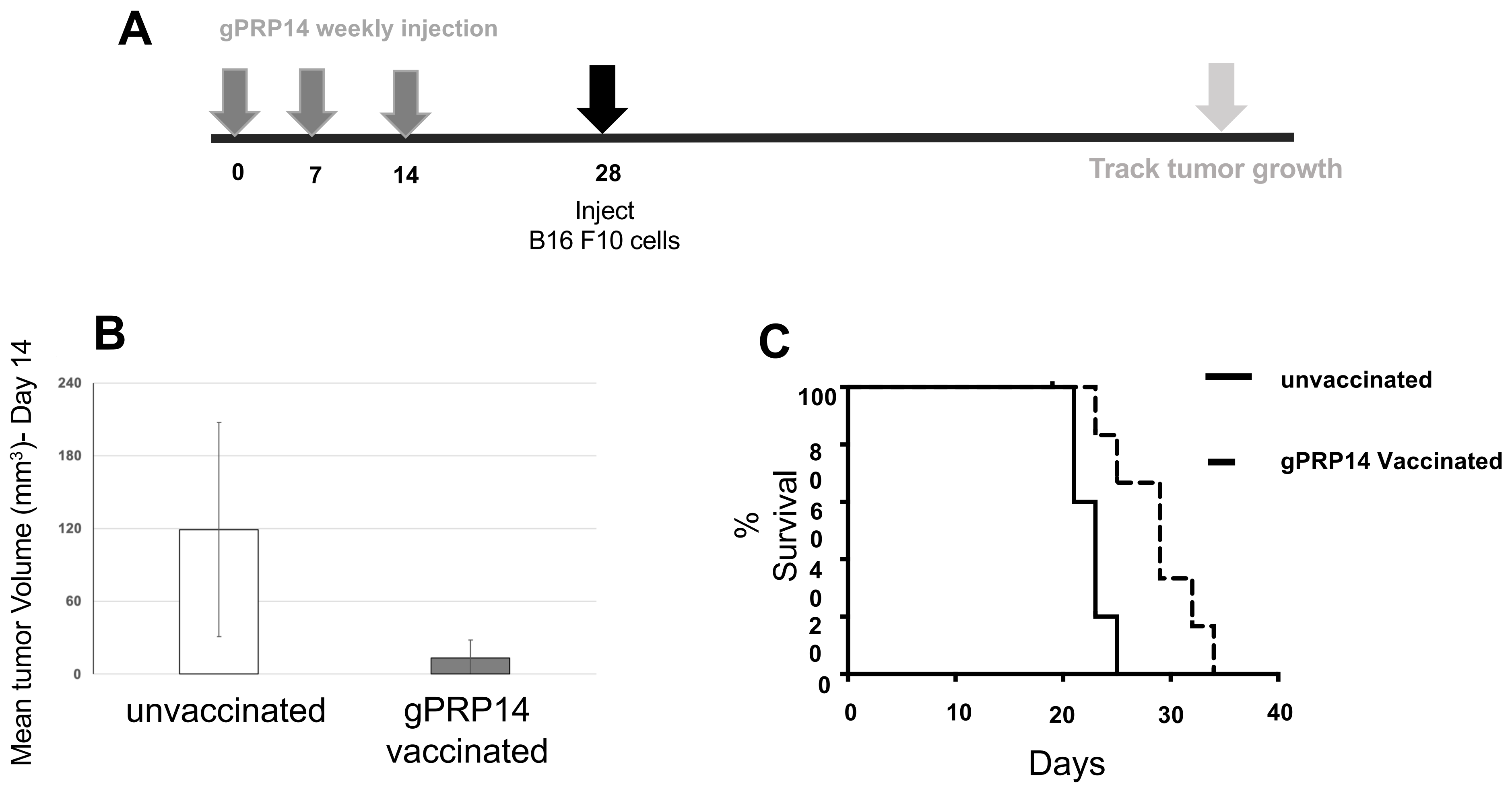


**Supp Fig 4 Low-dose vaccination with gPRP14 is partially protective in a melanoma syngeneic murine model. A)** Schedule of treatment of C57BL6 mice with gPRP14 (1 µg protein), followed by B16F10 melanoma cells inoculation. B) Mean tumor volume of B16F10 tumors in unvaccinated (White bar) or low-dose gPRP14 vaccinated (gray bar) mice, 14 days post tumor-implantation; C) Kaplan-Meier survival curves of unvaccinated (solid line) or low-dose gPRP14 vaccinated (dashed line) animals.

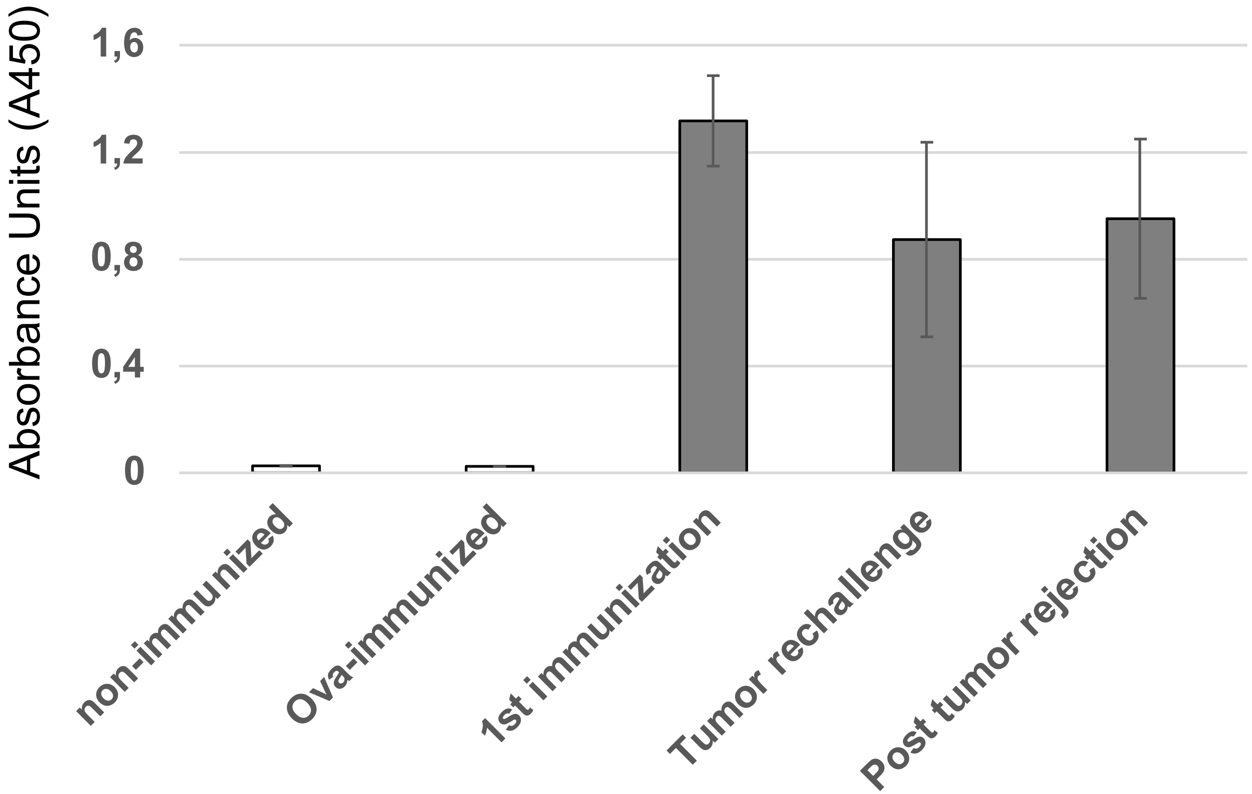


**Supp Fig 5. Mice maintain gPRP14 reactivity long-term post immunization during tumor re-challenge.**
ELISA measurement of gPRP14 serum reactivity :
-) at the time of initial B16F10 tumor cells implantation in control, ovalbumin-immunized and gPRP14-immunized mice (1st immunization)
-) in long-term survivor mice pre- (Tumor rechallenge) and post-secondary challenge with B16F10 melanoma cells (Post-tumor rejection).
